# Hippocampal damage causes retrograde amnesia for objects’ visual, but not odour, properties in male rats

**DOI:** 10.1101/2022.09.14.508050

**Authors:** Sean G. Lacoursiere, Brendan B. McAllister, Crystal Hadikin, Wayne W. Tschetter, Hugo Lehmann, Robert J. Sutherland

## Abstract

Damage to the hippocampus produces profound retrograde amnesia, but odour and object discrimination memories can be spared in the retrograde direction. Prior lesion studies testing retrograde amnesia for object/odour discriminations are problematic due to sparing of large parts of the hippocampus, which may support memory recall, and/or the presence of uncontrolled, distinctive odours that may support object discrimination. To address these issues, we used a simple object discrimination test to assess memory in male rats. Two visually distinct objects, paired with distinct odour cues, were presented. One object was associated with a reward. Following training, neurotoxic hippocampal lesions were made using *N-methyl-D-aspartate* (NMDA). The rats were then tested on the preoperatively learned object discrimination problem, with and without the availability of odour or visual cues during testing. The rats were also postoperatively trained on a new object discrimination problem. Lesion sizes ranged from 67-97% of the hippocampus (average of 87%). On the preoperatively learned discrimination problem, the rats with hippocampal lesions showed preserved object discrimination memory when tested in the dark (i.e., without visual cues) but not when the explicit odour cues were removed from the objects. Hippocampal lesions increased the number of trials required to reach criterion but did not prevent rats from solving the postoperatively learned discrimination problem. Our results support the idea that long-term memories for odours, unlike recall of visual properties of objects, does not depend on the hippocampus in rats, consistent with previous observations that hippocampal damage does not cause retrograde amnesia for odour memories.

## INTRODUCTION

Studies in humans and laboratory animals show that damage to the hippocampus produces profound retrograde amnesia. The magnitude of the loss of previously learned information is related to the extent of the damage to the hippocampus (Epp et al., 2008; Lehmann et al., 2010; Mumby et al., 1999; Sutherland et al., 2001). Hippocampal damage in male or female rats causes retrograde amnesia for a wide range of memory types, including place (Morris et al., 1982; Mumby et al., 1999), context (Sutherland et al., 2008; Broadbent & Clark, 2013), object recognition (Gaskin et al., 2003), shock-probe conditioning (Lehmann et al., 2006), and visual discrimination (Driscoll et al., 2005; Epp et al., 2008) memories. However, olfactory and object discrimination memories have been reported to be spared in the retrograde direction by similar hippocampal damage in male rats (Jonasson et al., 2004; Mumby et al., 1999). Olfaction is a highly conserved sensory system and may be the main sensory modality rodents use to interact with an environment (Ache & Young, 2005). The responses from different olfactory receptors are segregated in different cells in the olfactory epithelium, allowing discrimination between odorants (Ache & Young, 2005; Aqrabawi & Kim, 2020), perhaps explaining how the olfactory system in rodents can function independently of the hippocampus in storing and retrieving odour memories acquired during simple odour discrimination tasks.

Two-choice discrimination problems are commonly used to assess memory in rodents. The typical task involves presenting a rat with two cues, which could be objects, visual cues, odours, and so on, depending on the nature of the task. The rat is trained to criteria to select the rewarded cue (S+) and not the other (S-). In the testing phase, rats that have learned the task properly will persist in selecting S+ despite no longer receiving a reward. Hippocampal damage after visual discrimination training has been found to cause retrograde amnesia in male and female rats (Driscoll et al., 2005; Epp et al., 2008), suggesting the hippocampus is important for this type of memory. In odour discrimination tasks, however, rats with extensive hippocampal damage show postoperative performance similar to their preoperative performance. For example, in a study by Jonasson et al. (2004), male rats were presented with a forced decision on a simple odour discrimination. Following training, the rats were subjected to NMDA lesions and, after recovery, were tested on the preoperatively learned task. The rats with hippocampus damage showed equivalent recall to the intact rats. These results suggest that, unlike other types of memory, the memories used to solve odour discrimination problems may not be dependent on the hippocampus for recall.

Prior lesion studies conducted to test retrograde amnesia for object/odour discriminations are problematic because the objects used may have allowed for inadvertent use of odours to enable discrimination (Lehmann et al., 2007; Mumby et al., 1999; Wible et al., 1992), leading to ambiguity in the interpretation of the role of the hippocampus in object discrimination memory. Therefore, in the present study we trained male rats to discriminate between objects constructed of identical materials, to eliminate the use of intrinsic odour cues to differentiate between the objects. The objects were visually distinct and, during training, were paired with explicit odours that could be removed during testing, allowing the experimenter to control whether the rats could use odour cues to solve the problem.

Following nearly-complete hippocampal damage, the ability of the rats to solve the pre-operative object discrimination problem was tested with and without odour and visual cues. The rats showed preserved memory when the use of visual cues was eliminated, but retrograde amnesia for the same object discrimination problem when the explicit odour cues were removed. These results indicate that male rats with hippocampal lesions can use odour cues to solve a preoperatively acquired discrimination problem, supporting the idea that long-term memory for odours, unlike recall of visual cues, does not depend on hippocampal circuitry in rats.

## MATERIALS AND METHODS

### Subjects

The University of Lethbridge Animal Care Committee, which follows the guidelines set by the Canadian Council on Animal Care, approved all procedures. In total, 29 male Long-Evans hooded rats (Canadian Centre for Behavioural Neuroscience vivarium, Lethbridge, AB, Canada), weighing 250-350 g, were used. The rats were housed individually in standard laboratory cages on a 12:12 light-dark cycle (lights on at 7:00 AM), and all testing was done during the light phase. The rats had free access to water and were fed ∼25 g of rat chow per day throughout the experiment, except during the first week of recovery from surgery, during which food was given *ad libitum*.

### Apparatus

The apparatus for the object discrimination task has been described in detail elsewhere (Mumby et al., 1990). Briefly, it consisted of an elevated runway, separated from identical goal areas at each end by opaque guillotine doors. Each goal area contained two food wells, into which food pellets (45 mg Bio-Serv, Inc., Frenchtown, NJ, USA) could be delivered by hand through plastic tubes that were mounted on the outside of the apparatus. A short divider wall protruded from the center of the end wall and separated the two food wells.

The objects used as discriminanda (see Fig. 1) were large enough to cover a food well in the apparatus but small enough to be easily displaced by a rat. Specifically, the first pair of objects was a small house constructed out of red and blue Lego and a small step stool constructed of yellow and green Lego. The objects were of equal weight (25 g) and contained openings in which scented cotton balls could be placed. The openings were fashioned so that that the rats were unable to touch or lick the cotton balls. The second pair of objects was a plastic egg holder and a centrifuge tube (each weighing approximately 15 g). The objects were washed after each rat’s session with a solution of diluted chlorine bleach to remove any extraneous scents acquired during displacement by the rats or handling by the experimenter.

**Figure 1.**
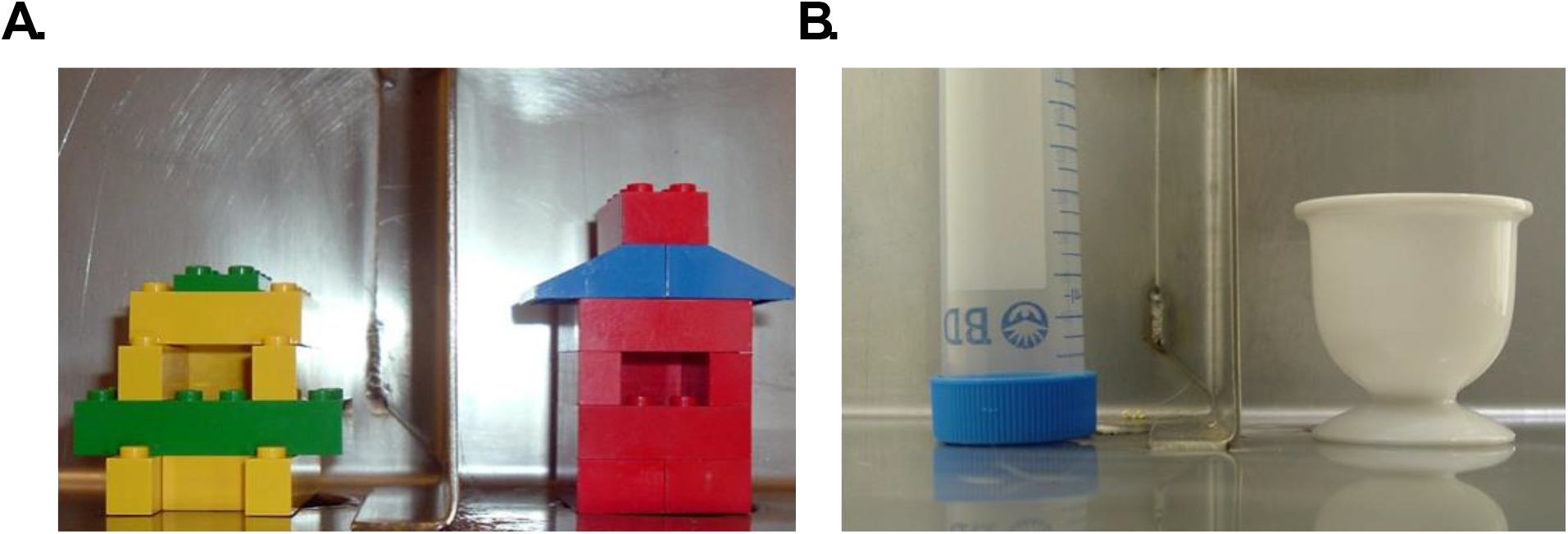
Discriminanda used for two object discrimination problems: Discrimination 1 (**A**), which was learned prior to hippocampal lesion surgery and tested postoperatively, and Discrimination 2 (**B**), which was learned and tested postoperatively.

The odours consisted of vanilla and peppermint concentrates manufactured by LorAnn Oils®. To present the odours, clean cotton balls containing 0.3 ml of oil were placed inside the objects each day. Based on previous reports (Scott et al., 2013), the odours used were chosen so that their subjective olfactory characteristics were distinct enough to discriminate between the two.

### Preoperative Object Discrimination Training

The timeline of the experiment is depicted in Figure 2. The rats were initially habituated to the apparatus and their behavior was shaped to retrieve food pellets from the food wells (see Mumby et al., 1990) across 4-7 daily sessions. At least 24 h after their last habituation/shaping session, the rats were trained on the first discrimination problem (Discrimination 1) in a single short session. The object pair consisted of the vanilla-scented Lego house and the peppermint-scented Lego stool (Fig. 1). One of the objects was designated S+ (rewarded) and the other was designated S- (not rewarded), counterbalanced within groups. The spatial position of S+ and S-was pseudorandomly counterbalanced across trials. Each rat was given a minimum of 40 trials, and the session ended when the rat reached criterion (7 consecutive choices of S+). The rats were then matched according to the number of trials they required to reach criterion and assigned to either the sham lesion (SHAM) or hippocampal lesion (HPC) group.

**Figure 2.**
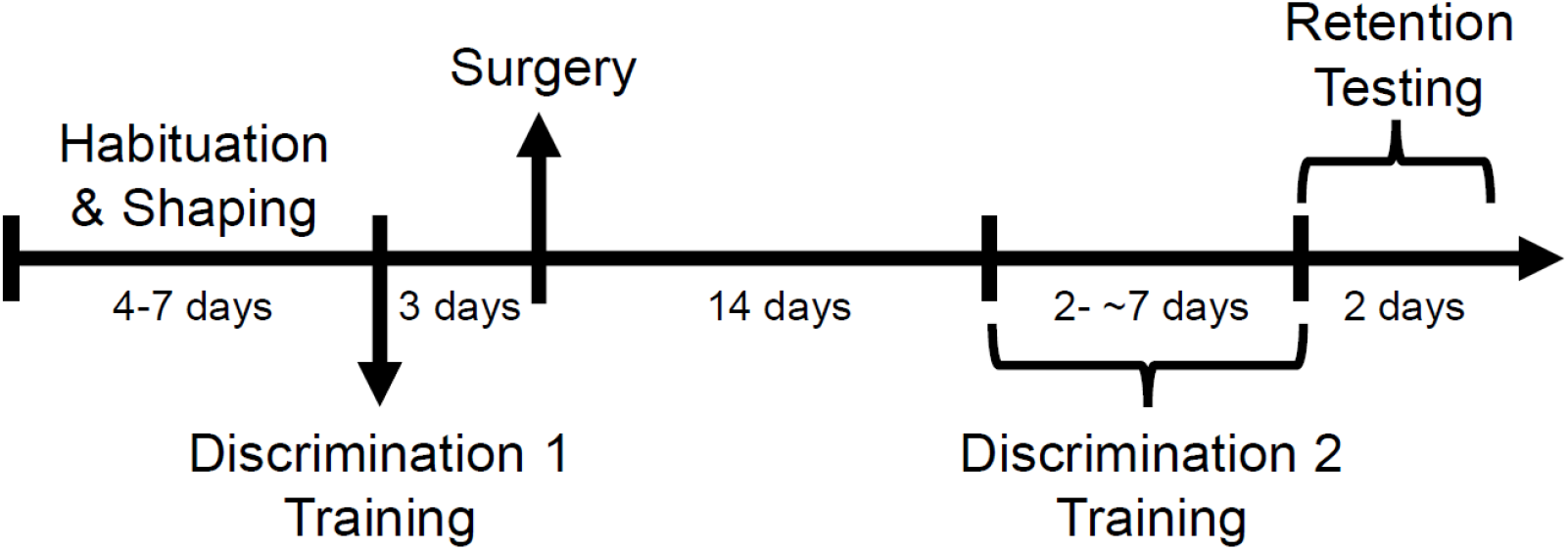
Timeline diagram of the experiment. After habituation to the apparatus and behavioural shaping, the rats were trained on an object discrimination (Discrimination 1) with explicit odours in a single learning session. Three days later, the rats received sham or hippocampal lesion surgery (Surgery) followed by a 14-day surgical recovery period. The rats were then trained on a second object discrimination (Discrimination 2) over a few days until they reached a success criterion of 80%. Finally, they were tested for retention of Discrimination 1 with intermixed trials from Discrimination 2. Note that the rats were tested for two days, and Discrimination 1 explicit odours or visual cues were removed on one of these days.

### Surgery

Approximately 72 h after training on Discrimination 1, the rats underwent surgery. Hippocampal lesions were completed following the protocol described by Lehmann et al. (2010). Briefly, the rats were anaesthetized with isoflurane (Janssen, Toronto, ON, Canada) in 1.0 L/min oxygen at 21 °C (Benson Medical Industries, Markham, ON, Canada) and administered buprenorphine (0.07 cc, 0.3 mg/mL, i.p.; Reckitt & Colman, Richmond, VA, USA) as an analgesic. The rats were then transferred to a stereotaxic frame (Kopf Instruments, Tujunga, CA, USA), and the scalp was cleaned with 4% stanhexidine followed by 70% isopropyl alcohol. A midline incision was made to expose the skull, and the periosteum was excised. The rats in the HPC group received a complete bilateral lesion of the hippocampal formation made by intrahippocampal injections of NMDA (7.5 μg/μL in 0.9% saline; Sigma Chemical Co., St. Louis, MO, USA) at ten sites bilaterally, following coordinates from Lehmann et al. (2010). The infusions were done sequentially through a 30-gauge injection cannula attached to a 10 μL Hamilton syringe by polyethylene tubing (PE-50). At each site, 0.4 μL was infused at a rate of 0.15 μL/min. The cannula was left in place for 2.5 min following injection to facilitate diffusion. The scalp was closed using wound clips following the injections. Diazepam (0.2-0.6 cc, 10 mg/mL, i.p.; Sabex, Boucherville, QC, Canada) was given to the rats as a prophylactic against seizures as they awakened. The same surgical procedure was given to the rats in the SHAM group, but no damage was done to the skull or brain of these rats. This sham surgical procedure (not drilling into the skull and/or not lowering the infusion cannula into the cortex overlying the hippocampus) is now common across many labs investigating the effects of hippocampal damage on memory, including retrograde memory (Broadbent et al., 2007; Epp et al., 2008; Gaskin et al., 2003; Gidyk et al., 2021; McDonald et al., 2018; Quinn et al., 2008; Tse et al., 2007; Winocur et al., 2005, 2013).

### Postoperative object discrimination training

Fourteen days after surgery, the rats were trained on a new object discrimination problem (Discrimination 2). The purpose of Discrimination 2 was, first, to motivate the rats to continue displacing the objects on the Discrimination 1 extinction trials during retention testing and, second, to correct any locomotor abnormalities/thigmotaxis in the object discrimination performance that may have resulted from the lesions (Day et al., 1999; Coutureau et al., 2000; Bannerman et al., 2001). Training the rats postoperatively on a second discrimination also allowed for assessment of the effects of the lesion on object discrimination acquisition, and it provided a way to distinguish the anterograde and retrograde effects of the lesions on memory.

For Discrimination 2, a new pair of objects (egg holder and centrifuge tube), with no odour added to either object, served as the discriminanda (Fig. 1). As before, one of the objects was designated S+ and the other one was designated S-, counterbalanced within groups. The rats were given 20 training trials per day for a minimum of 2 days. The training continued for a given rat until it reached a criterion of at least 16 out of 20 choices of S+ in a session.

### Object Discrimination Retention Testing

The rats’ retention for the object discrimination problems was assessed 24 h after criterion was reached for Discrimination 2. The rats were tested over 2 days to determine the retention of Discrimination 1. On one day, the rats were tested for Discrimination 1 with the explicit peppermint and vanilla odour cues and under the original visual/lighting conditions (Standard test). On the other day, the rats assigned to the No-Odour condition were tested without explicit odour cues (i.e., with the peppermint and vanilla odour cues removed), increasing the need for the rats to rely on the visual cues to succeed on the discrimination test. In contrast, the rats assigned to the Dark condition were tested in complete darkness. In this instance, the rats needed to rely on olfactory cues to complete the discrimination, and the experimenter used night vision goggles to conduct the testing.

The order of the testing days (Standard vs. No-Odour or Dark) was counterbalanced, as was the order in which the rats were tested. For each testing condition, the rats were given 2 trials of Discrimination 1, and their selections of S+ were not rewarded (i.e., extinction trials). These trials were then followed by 2 trials of Discrimination 2, in which selection of S+ was rewarded. This sequence was continued until 10 trials for each problem were completed. Subjecting rats to extinction trials for Discrimination 1 allowed confidence that any selection bias in the retention test could be attributed to the rat’s memory of the preoperative acquisition, with no influence from post-surgery reinforcement. The interleaving of the rewarded Discrimination 2 trials served to maintain the rat’s motivation to displace the objects.

### Histology

Upon completing the behavioural testing, the rats were given an overdose of sodium pentobarbital (0.5 cc; 320 mg/ml, i.p.; Schering Canada Inc., Pointe-Claire, QC, Canada) and intracardially perfused with 0.9% saline followed by 10% formalin. Their brains were collected and stored in 10% formalin-30% sucrose for at least 48 h. The brains were sectioned at 40 μm thickness using a freezing sliding microtome. Every twelfth section of the hippocampus was mounted on gelatin-coated glass slides, and the sections were stained with Cresyl violet and cover slipped. Coronal sections spanning the entire hippocampus were imaged and digitized using a slide scanner (NanoZoomer-RS, Hamamatsu).

Hippocampal damage was estimated using principles of the Cavalieri method, following the protocol described by Scott et al. (2016), with minor modifications. A systematic sampling grid (with an area per square of 0.0144 mm^2^) was randomly laid over each image, and the number of points hitting the dentate gyrus granule cell layer or the CA1/CA3 pyramidal cell layers was counted. Grids were generated using ImageJ software (https://imagej.nih.gov/ij). For each HPC rat, the total number of hits was divided by the average number of hits obtained from three SHAM rats (i.e., 1209 hits), resulting in the proportion of remaining hippocampal tissue. The complement proportion was used as an estimate of the percentage of hippocampal damage.

### Statistics

SPSS Statistics (Version 27, IBM) and GraphPad Prism 8 (Version 8.4.3; macOS) were used for statistical analysis and figure production, respectively. An alpha level of 0.05 was used. To test whether the rate of S+ responses was significantly greater than would be predicted by chance (i.e., greater than 50%) within each group, one-tailed one-sample *t*-tests were used. Comparisons between two groups were made using two-tailed independent-samples *t*-tests. A two-way analysis of variance (ANOVA) was used to determine if S+ responses in the Discrimination 1 retention test were affected by lesion (SHAM vs. HPC) or testing condition (Standard vs. No-Odour vs. Dark). Equal variances were not assumed if Levene’s test was significant. Means reported in the text below are presented ± the standard error of the mean (SEM).

## RESULTS

### Hippocampal Lesions

The NMDA injections produced extensive cell loss across all principal subfields of the hippocampus (see Fig. 3 for a representative lesion). Estimates of hippocampal damage in the HPC rats from the No-Odour condition (*n* = 7) ranged from 84%-95% with a mean of 90% ± 2%. Estimates of hippocampal damage in the HPC rats from the Dark condition (*n* = 7) ranged from 67%-97%, with a mean of 87% ± 5%. The dorsal hippocampus was completely damaged in all the HPC rats, and any observed sparing of the hippocampus was predominantly located in the most anterior or posterior parts of the ventral hippocampus. Damage to the overlying neocortex was minimal across the HPC rats and limited to the sites where the injection cannula was lowered. Overall, the average lesion sizes observed were comparable to, or slightly larger than, the average lesion sizes reported in other object, odour, and visual discrimination or recognition studies using the same lesion method and number of injection sites (Jonasson et al., 2004; Driscoll et al., 2005; Mumby et al., 2005; Lehmann et al., 2007).

**Figure 3.**
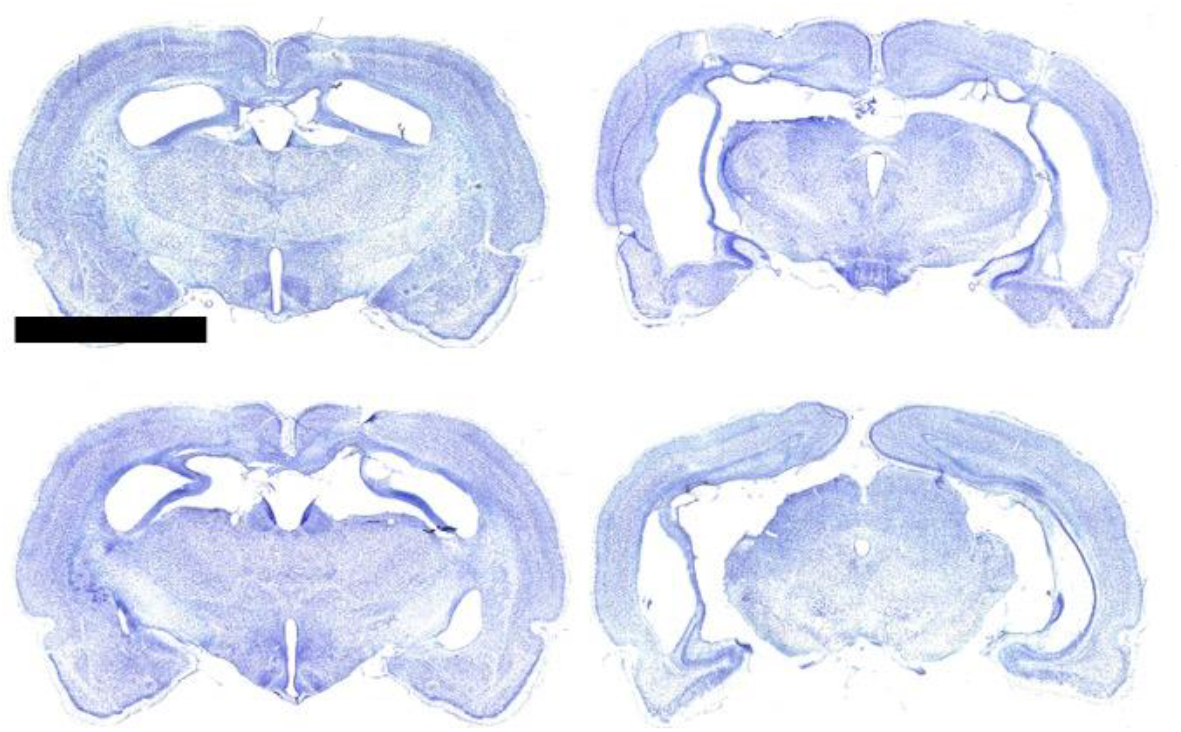
Photomicrographs of Nissl-stained sections from a rat brain with a representative lesion, estimated at 88% of the hippocampus. Scale bar represents 5 mm.

### Object Discrimination

#### Discrimination 1 – Preoperative training

The SHAM and HPC rats, whether from the No-Odour or Dark condition, learned the preoperative object discrimination problem (Discrimination 1) at a similar rate. The SHAM No-Odour group (*n* = 8) required 96.4 ± 10.6 trials to reach criterion, the SHAM Dark group (*n* = 7) required 88.0 ± 8.7 trials, the HPC No-Odour group (*n* = 7) required 99.9 ± 13.6 trials, and the HPC Dark group (*n* = 7) required 90.0 ± 10.9 trials. A two-way ANOVA revealed no main effect of lesion (SHAM vs. HPC) [*F*_1,25_ = 0.06, *p* = 0.807], no main effect of cue (No-Odour vs. Dark) [*F*_1,25_ = 0.68, *p* = 0.418], and no interaction [*F*_1,25_ < 0.01, *p* = 0.947]. Moreover, the time required to learn the discrimination did not differ across the lesion groups [*F*_1,23_ < 0.01, *p* = 0.951] or cue [*F*_1,23_ = 0.43, *p* = 0.516], with the group means ranging from 51-63 min. Two time values were missing due to a recording problem – one from the SHAM No-Odour group and one from the HPC Dark group – but the missing times were not noted to be out of range. These results confirm that the rats assigned to each group were properly matched for their acquisition performance.

#### Discrimination 1 - Postoperative retention

Figure 4 illustrates the retention performance of the SHAM and HPC rats on Discrimination 1 when tested in the Standard (visual and odour cues present), No-Odour (discrete odour cues removed), and Dark (complete darkness) conditions. The data from the two SHAM groups were pooled for the Standard test because they did not statistically differ [77.5% vs. 74.3% S+ choices; *t*_13_ = 0.27, *p* = 0.789]. The same was done for the data from the two HPC groups on the Standard test, again because of similar performance [81.4% vs. 67.1% S+ choices; t_12_ = 1.01, *p* = 0.333]. This pooling simplified presentation of the data and overall analysis.

**Figure 4.**
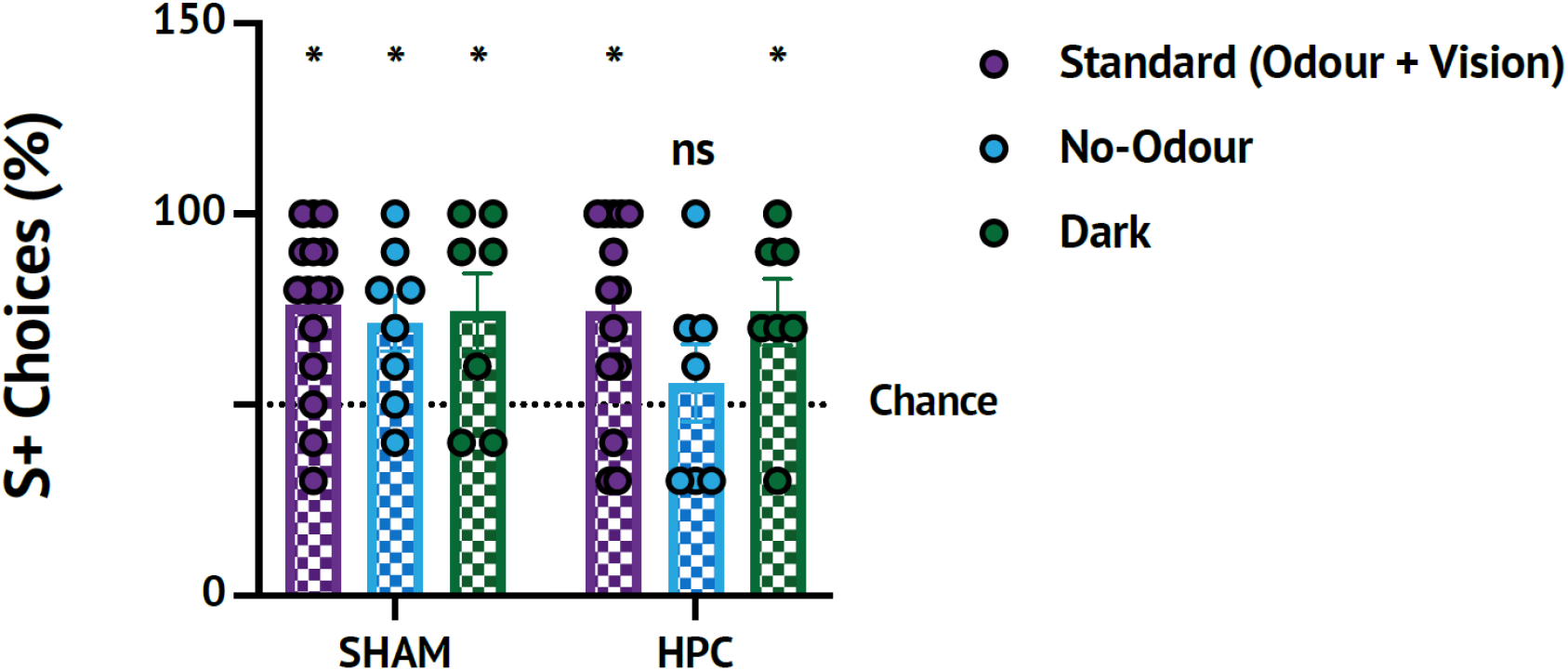
Postoperative performance of rats on Discrimination 1, tested with or without the explicit odour or visual cues that were present during preoperative acquisition of the problem. For each group, performance was assessed by comparing the rate of S+ selections to the rate predicated by chance (i.e., 50%, dotted line on figure). Whereas the SHAM rats could solve the problem with odour and visual cues intact (Standard; *n* = 15), without the explicit odour cues (No-Odour; *n* = 8), and without visual cues (Dark; *n* = 7), the HPC rats were impaired in the No-Odour condition (*n* = 7), but not in the Standard (*n* = 14) or Dark condition (*n* = 7). Error bars = SEM

In the postoperative retention test for Discrimination 1 (Fig. 4), a two-way ANOVA showed that there was no significant main effect of testing condition [*F*_2,52_ = 1.22, *p* = 0.303] or lesion [*F*_1,52_ = 0.73, *p* = 0.397], nor was there an interaction between the two factors [*F*_2,52_ = 0.49, *p* = 0.615]. Comparing the performance of the rats in each condition to chance, one-sample *t*-tests showed that the SHAM rats could solve the problem under all the conditions, as indicated by selecting S+ at a rate significantly greater than 50% [Standard: *t*_14_ = 4.58, *p* < 0.001; No-Odour: *t*_7_ = 2.96, *p* = 0.011; Dark: *t*_6_ = 2.38, *p* = 0.027]. In contrast, the HPC rats performed no better than chance under the No-Odour condition, selecting S+ on 55.7 ± 10.2% of the trials [*t*_6_ = 0.56, *p* = 0.298]. However, the HPC rats were able to select S+ at a rate significantly greater than chance when they could rely on odour cues, both in the Standard and Dark conditions [Standard: *t*_13_ = 3.43, *p* = 0.002; Dark: *t*_6_ = 2.80, *p* = 0.016].

#### Discrimination 2 – Postoperative training and retention

Following surgery and recovery, the SHAM and HPC rats were trained to criterion on a new discrimination problem (Discrimination 2). The HPC rats required significantly more trials to reach criterion than did the SHAM rats [*t*_27_ = 2.83, *p* = 0.009; Fig. 5A]. Despite this difference in acquisition, the performance of the SHAM and HPC rats on the final training session, as assessed by the percentage of S+ selections out of the 20 trials, was very similar [*t*_27_ = 0.34, *p* = 0.735; Fig. 5B]. Furthermore, on the Discrimination 2 retention test, performance did not significantly differ between the HPC and SHAM rats [*t*_27_ = 1.87, *p* = 0.072; Fig. 5C]. These results show that, while the HPC rats required more trials to learn the anterograde discrimination task compared to the SHAM rats, by the end of training and during retention testing the SHAM and HPC rats showed similar performance.

**Figure 5.**
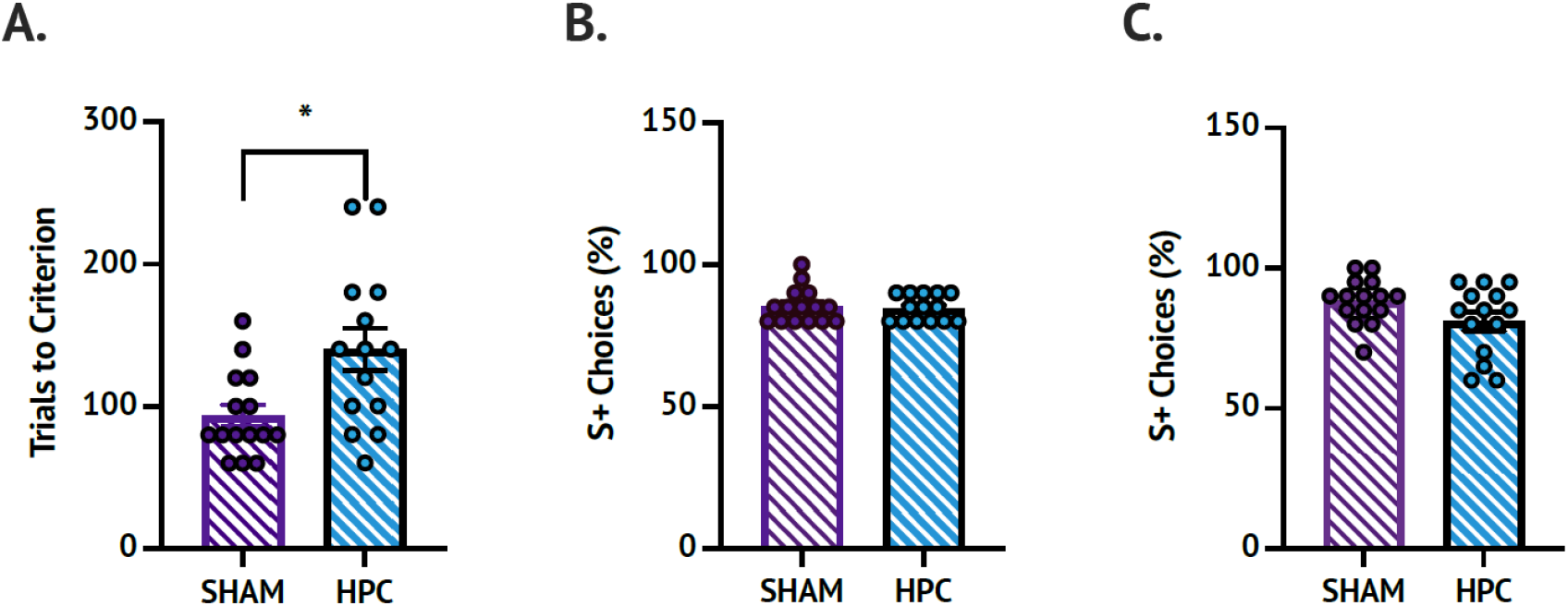
Performance of rats on an object discrimination problem (Discrimination 2) acquired after the rats had received either SHAM (*n* = 15) or HPC (*n* = 14) lesions. **A**. Trials to criterion during acquisition of Discrimination 2. The HPC rats required significantly more acquisition trials to reach criterion than did the SHAM rats. **B.** The S+ choice rate during the final session of Discrimination 2 acquisition was similar between the SHAM and HPC rats. **C.** The S+ choice rate during Discrimination 2 retention testing did not differ significantly between groups. Error bars = SEM.

## DISCUSSION

The purpose of this study was to test whether memory for discrimination problems acquired before receiving a complete, bilateral hippocampal lesion is affected by the presence or absence of odour and visual cues. To this end, rats were trained on an object discrimination problem (Discrimination 1), which could be solved using visual cues, explicit odour cues (i.e., strong, distinct odours applied to the objects by the experimenter), or tactile cues. In principle, the use of implicit odour cues to solve the problem is a possibility, but it is unlikely in the present experiment, because the objects were constructed out of identical materials and so would very likely not differ in scent. The Discrimination 1 problem was successfully acquired by the rats in a single, short training session. Retention for the problem was then tested after hippocampal lesions. In one condition (No-Odour), retention was tested without the explicit odour cues that were associated with the objects during acquisition. In another condition (Dark), the contribution of visual cues was assessed by testing the rats’ retention performance with the lights off and no other light source present, making the room completely dark.

Our key finding was that NMDA lesions encompassing almost the entirety of the hippocampus did not cause retrograde amnesia for the object discrimination problem when rats could use odour cues to make a choice. This was the case even when the rats were tested in the dark, rendering them unable to use visual cues. With the explicit odours present in the retention test, the HPC rats selected S+ at a rate significantly greater than would be predicted by chance. However, when the explicit odour cues were removed during the retention test, forcing the rats to rely on visual or other cues to solve the problem, the HPC rats failed to select S+ at a rate greater than chance, suggesting retrograde amnesia for the object/reward association. The SHAM rats, in contrast, were able to solve the problem without using the explicit odour cues. It is unlikely that visual impairments caused by hippocampal damage contributed to the failure of the HPC rats to solve the discrimination problem in the No-odour condition, because male and female rats with similar hippocampal lesions were not impaired at learning simple visual discriminations in a visual-only task (Driscoll et al., 2005; Epp et al., 2008).

Our results suggest that there may be something unique about how the brain stores odour memories, relative to memories derived from vision and perhaps other sensory modalities. Whatever these unique neurobiological features may be, they allowed the odour features of an object/reward association memory, learned with an intact hippocampus, to be successfully recalled without the hippocampus. In contrast, memory for the non-odour features (e.g., colour and shape of the objects) did not survive a complete lesion of the hippocampus. This idea is consistent with previous findings that complete hippocampal lesions cause retrograde object recognition memory deficits when male rats are tested in the novel object preference test using objects without explicit odours (Broadbent et al., 2010; Gaskin et al., 2003). It is also consistent with the observation in male and female rats that hippocampal lesions cause retrograde amnesia for the association between a visual cue (e.g., a picture) and an escape platform in a visual discrimination test (Driscoll et al., 2005; Epp et al., 2008; Lehmann et al., 2021; Sutherland et al., 2001). Finally, it is consistent with previous findings in male rats that hippocampal lesions do not cause retrograde amnesia for odour recognition memories (Scott et al., 2013) or in an odour discrimination task (Broadbent et al., 2007).

In several studies, male rats with complete hippocampal lesions did not show retrograde amnesia in object discrimination tasks similar to the one used in the present study (Lehmann et al., 2007; Mumby et al., 1999; Wible et al., 1992). In another study, male rats with hippocampal lesions performed worse than control rats, but did not show complete retrograde amnesia, as evidenced by the ability to solve the discrimination problem at a rate greater than chance (Broadbent et al., 2007). Importantly, none of these previous studies reported pairing objects with explicit odour cues. According to the interpretation presented above, without explicit odour cues, the rats in these studies should not have been able remember the object/reward association after hippocampal damage. An alternate explanation is required, then, that addresses the question of why the HPC rats in the present study failed to recall the object/reward association when they could not rely on explicit odour cues, whereas rats in previous studies that did not use explicit odour cues performed the task adequately. One possibility is that, in at least some previous studies, the rats with hippocampal lesions may not have needed explicit odour cues because they could rely on implicit ones instead. In the study by Wible et al. (1992), objects made of varying materials (e.g., metal, rubber) were used as discriminanda, potentially making it so that the rats could discriminate between the objects based on odour. In other studies, the authors reported using objects with no obvious scents (Mumby et al., 1999) or consistently used objects made of plastic (Broadbent et al., 2007; Lehmann et al., 2007). However, in these studies it was not reported that objects made of identical materials were used, as was done for Discrimination 1 in the present study to minimize the ability of rats to use implicit odour cues to solve the discrimination problem. Notably, the postoperatively acquired Discrimination 2 used two different plastic objects, and the rats were able to discriminate between them in complete darkness, though whether the rats relied on implicit odour cues, tactile cues, or both to solve this problem is not known.

Cue overshadowing may offer another explanation. The rats trained on the discrimination problem without explicit odour cues are, by default, forced to rely on other information, such as visual and tactile cues, to solve the problem. The rats therefore pay close attention to these cues, allowing the rats to form a strong memory of the object that is resilient to hippocampal damage. When rats are trained on the same task *with* explicit odour cues, the rats may rely heavily on these odours to solve the problem – perhaps because odour cues are particularly salient to a rat – and pay less attention to the non-odour features of the object. This might result in a weaker memory of these features, which is sufficiently strong for a normal rat to recall the object/reward association, but not strong enough for a rat without a hippocampus to do so. In other words, it is not that rats with hippocampal lesions are incapable of recalling object/reward associations formed pre-lesion, but rather that the explicit odour cues overshadow the non-odour cues, preventing the rats from forming a hippocampal-independent memory of the object/reward association that they *would* have been able to form had the explicit odours not been present. Because the objects used as discriminanda in the present study were compound stimuli with both olfactory and visual features, the possibility of cue overshadowing cannot be excluded. This limitation of using compound stimuli is offset by the advantage that it ensures the animals’ history of contact and reward with the olfactory and visual cues is as similar as possible during acquisition.

Regardless of which interpretation is correct, the totality of the evidence suggests that odour memories are less susceptible to hippocampal damage than are similar memories involving non-odour information. Previous studies have shown that complete hippocampal lesions do not affect the ability to form new odour recognition memories and maintain these memories over long (i.e., 5 week) retention intervals, nor do lesions disrupt the ability to recall odour recognition memories acquired preoperatively (Feinberg et al., 2012; Hunsaker et al., 2008; Scott et al., 2013). Similarly, hippocampal damage in male rats does not cause retrograde or anterograde amnesia for odour/reward associations in odour discrimination memory tasks (Broadbent et al., 2007; Jonasson et al., 2004; Wood et al., 2004). These findings, in conjunction with the present results, indicate that extrahippocampal systems in the brain are responsible for the long-term maintenance of odour memories. The striatal memory system may be particularly important for remembering odour/reward associations, as lesions of dorsal striatum in male rats have been found to cause retrograde amnesia and impair, though not abolish, memory in the anterograde direction on an odour discrimination task (Broadbent et al., 2007).

This is not to say that the hippocampus has no role in odour processing or in odour memory tasks. For instance, lesioning the ventral dentate gyrus with colchicine in male rats impairs performance in a delayed-matching-to-sample odour discrimination working memory test (Weeden et al., 2014). The hippocampal damage did not impair ability to differentiate between odours, as indicated by normal performance when the delay between sample and test was short (15 s). However, at a longer delay (60 s), the rats with lesions were impaired, particularly when they were required to distinguish between similar odours. With larger lesions to the ventral hippocampus, male rats were impaired even at the 15 s delay, though they could still discriminate between odours in a simple discrimination task (Kesner et al., 2011). Moreover, some studies have shown retrograde amnesia for odour/flavour memories acquired very shortly (i.e., a couple of days) before hippocampal lesions are sustained in male rats (Tse et al., 2007; Winocur, 1990; Winocur et al., 2001), suggesting that odour recognition memories may be transiently dependent on the hippocampus. The possibility should not be dismissed, however, that this short period of retrograde amnesia may result from disrupted cellular consolidation in structures other than the hippocampus, as lesions targeted to the hippocampus may temporarily disrupt cellular consolidation even in off-target structures (Glenn et al., 2005; Rudy & Sutherland, 2008). In any case, none of these studies contradicts the idea that long-term storage of odour memories, particularly relatively simple odour recognition or odour discrimination memories, is independent of the hippocampus.

The story is similar for other brain structures directly upstream of the hippocampus, in that the bulk of the evidence indicates that these structures are involved in performing certain odour-based tasks, but each structure on its own is not essential for simple odour memories. The lateral entorhinal cortex (LEC) is a primary source of input to the hippocampus and receives projections from olfactory areas like the olfactory bulbs and the piriform cortex (Kerr et al., 2007). Reversibly inactivating the LEC impairs the performance of male rats in an odour discrimination task when a fine discrimination is required, demonstrating a role for the LEC in processing odour information (Chapuis et al., 2013). The task used by Chapuis and colleagues involved making an immediate response after sampling the odours, and so it does not reveal much about the role of LEC in odour memory; but, aspiration of the LEC in male rats does not impair the ability to form new odour recognition memories or to retain these memories for intervals up to 48 h (Wirth et al., 1998). Transection of the lateral olfactory tract - a major source of olfactory input to the entorhinal cortex and, therefore, also to the hippocampus - had no effect on odour/reward association memory in an odour discrimination task in male rats (Slotnick & Risser, 1990). However, when this transection was combined with electrolytic lesions of the mediodorsal thalamic nucleus, part of the olfactory thalamocortical system, rats did show anterograde amnesia in this task. In the case of the perirhinal cortex, which also receives input from the piriform cortex and projects directly and indirectly to the hippocampus (Furtak et al., 2007), excitotoxic lesions cause anterograde impairments of recognition memory for social odours at relatively long retention intervals (1-48 h), but not of recognition memory for more distinct non-social odours (household spices) at the same intervals in male rats (Feinberg et al., 2012).

A secondary finding of the present study was that complete hippocampal lesions do not prevent learning and memory of new object/reward associations, nor do they impair the ability to discriminate between distinct objects. This was observed when rats were tested on the Discrimination 2 problem, which was acquired post-lesion. Hippocampal lesions may have had some effect on the ability to learn and remember Discrimination 2. Specifically, relative to the SHAM rats, the HPC rats required more trials to learn this discrimination. However, with additional training, the HPC rats did eventually reach a level of performance that was very similar to the level achieved by the SHAM rats. Moreover, on the Discrimination 2 retention test, the HPC rats did not perform significantly worse than the SHAM rats, and both groups performed well above chance levels. Thus, complete hippocampal lesions may cause some mild anterograde object discrimination learning impairments, but clearly do not cause complete anterograde amnesia for object/reward association memories. This is consistent with previous findings that male rats with hippocampal lesions are still able to form new object recognition memories, as assessed using the novel object preference test (Forwood et al., 2005; Gaskin et al., 2003; Mumby et al., 2002, 2005), and new object/reward associations, as assessed using object discrimination tasks (Broadbent et al., 2007; Lehmann et al., 2007; Mumby et al., 1999). Male rats with hippocampal damage have been shown to exhibit small decreases in novel object preference relative to sham-lesioned animals, but object recognition memory was still intact in these lesioned animals, as indicated by preference for the novel object significantly greater than chance (Broadbent et al., 2010). Based on our present findings, and on previous work using the same object discrimination apparatus (Lehmann et al., 2007), there is no reason to believe that learning Discrimination 2 interfered with Discrimination 1.

The design of the present study included only male rats, limiting any conclusion about potential sex differences. There is considerable evidence suggesting that male and female rats may use different spatial navigation strategies (Koss & Frick, 2017; Yagi & Galea, 2018), but little evidence for sex differences in recognition or discrimination memory (Becegato & Silva, 2022). Also, hippocampal lesions similarly affect male and female rats across most memory tasks (Morris et al., 1982; Mumby et al., 1999; Driscoll et al., 2005; Epp et al., 2008; Sutherland et al., 2008; Broadbent & Clark, 2013). Hence, we speculate that our observed findings would likely replicate in female rats.

In summary, we observed that complete hippocampal lesions in rats caused retrograde, but not anterograde, amnesia in an object discrimination task, so long as the task could not be solved using odour cues. When the rats were able to use odour cues to solve the discrimination problem, they were able to perform the task successfully, demonstrating that memory for the odour features of an object/reward association memory is spared following hippocampal damage. When considered alongside previous findings, our results support the idea that object/reward association memories that would normally be hippocampal-independent can be made dependent by pairing the objects with explicit odour cues during acquisition and then removing these odour cues during retention testing.

## Acknowledgements

The authors thank Valérie Lapointe for assistance with histology. Funding was provided by National Sciences and Engineering Research Council of Canada (NSERC) Discovery Grants to HL and RJS.

## Abbreviations

ANOVA: analysis of variance
LEC: lateral entorhinal cortex
NMDA: N-methyl-D-aspartate
SEM: standard error of the mean.

